# IL-36γ signalling promotes a proinflammatory macrophage state that is associated with reduced lipid uptake

**DOI:** 10.1101/2025.10.22.683795

**Authors:** Jillian Yong Xin Sieh, Gabriel Osborn, Sophia N. Karagiannis, Francesca Capon

**Author notes:** Corresponding authors: Sophia Karagiannis, St John’s Institute of Dermatology, School of Basic & Medical Biosciences, King’s College London, 9^th^ Floor Tower Wing, Guy’s Hospital, Great Maze Pond, London SE1 9RT, UK; Francesca Capon, Centre for Molecular Medicine and Therapeutics 950 West 28th Avenue, Vancouver, BC, Canada V5Z 4H4.

## Abstract

Macrophages exhibit remarkable functional plasticity, adopting immune activating or immune suppressive states in response to environmental cues. While these phenotypic shifts are essential to immune homeostasis, the mechanisms whereby they are regulated in humans are poorly understood. Here, we investigated the role of interleukin (IL)-36γ, a barrier cytokine that is strongly induced in response to infection. We show that IL-36γ signalling modifies the anti-inflammatory phenotype of human alternatively activated (M2a) macrophages, by decreasing the expression the CD163 M2 marker. This change was accompanied by the upregulation of M1 surface markers (CD40, CD80) and M1 cytokines (e.g. TNFα, CXCL8). IL-36γ treatment of M2a macrophages also reduced the expression of TREM2 and CD36, two lipid-binding receptors that sustain the energy requirements of the M2 state. Accordingly, we also observed a decrease in the uptake of CD36 and TREM2 ligands (oxidised low-density lipoproteins). These findings indicate that IL-36γ shifts the immune and metabolic profile of M2a macrophages towards an M1-like state.

## 1. Introduction

Macrophages display remarkable functional plasticity and respond to environmental cues by adopting a variety of activation states. To reflect this diversity, macrophages are broadly classified into immune activating and immune suppressive populations. In-vitro, these are studied as M1 and M2 subsets. M1 macrophages have immune activating functions mediated by the production of inflammatory cytokines and the expression of T-cell co-stimulatory molecules. M2 macrophages can be further subdivided in M2a, M2b, M2c and M2d subsets, respectively contributing to tissue repair, immune regulation, resolution of inflammatory responses and tumour progression^1^. The balance between immune activating and immune suppressive populations is critical to tissue homeostasis, as demonstrated by the dysregulation of macrophage states in chronic inflammatory disease and cancer^1^. While targeting macrophage polarisation has emerged as a new strategy for the treatment of these conditions, the factors that promote macrophage phenotype shifts in humans are not fully understood.

Interleukin (IL)-36α, IL-36β, and IL-36γ (IL-36) are closely related members of the IL-1 cytokine family. They bind a common receptor that consists of a signalling subunit (IL-36R) and an accessory co-receptor (IL-1RAcP). IL-36 is primarily expressed in skin, lung, and gut, where it is rapidly induced following microbial challenge or tissue injury. Engagement of IL-36R then triggers canonical NF-κB and MAPK signalling, resulting in the release of pro-inflammatory cytokines and chemokines, which recruit and activate myeloid and lymphoid cells^2^.

IL-36 amplifies innate and adaptive immune responses. In the skin, it promotes keratinocyte activation and antiviral responses^3^. In the gut, it strengthens mucosal defences by upregulating antimicrobial peptide production^4^. Finally, IL-36 enhances responses to infection and inflammatory stimuli in the lung, while also contributing to tissue injury^5^. In all barrier sites, IL-36 upregulates dendritic cell activity, thus promoting the polarisation of T cells towards Th1 and Th17 phenotypes^6,7^.

Given these proinflammatory functions, we hypothesised that IL-36 can shift M2 macrophages towards an M1-like state. We have therefore explored the effects of the cytokine on macrophage surface markers and cytokine production.

## 2. Materials and Methods

### 2.1 Generation and Stimulation of Human Monocyte-Derived Macrophages

Leukocyte-enriched cones obtained from healthy donors were purchased from the NHS Blood and Transplant Service (Tooting, UK) and Peripheral blood mononuclear cells (PBMCs) were isolated using Ficoll-Paque™ PLUS (Cytiva). Monocytes were then extracted from freshly isolated PBMCs with the Pan Monocyte Isolation Kit (Miltenyi Biotec). Macrophages were differentiated and polarised as previously described^8^. Briefly, monocytes were grown in complete medium (RPMI 1640-GlutaMAX™ supplemented with 10% FBS and 1% penicillin-streptomycin) containing 50 ng/mL M-CSF for four or five days. Cells were then polarised by adding complete medium supplemented with 16.7 ng/mL M-CSF and with the agents required for M1 (100 ng/mL LPS (Merck) + 40 ng/mL IFN-γ (Thermo Fisher Scientific)), M2a (20 ng/mL IL-4 (PeproTech)) or M2c (20 ng/mL IL-10 (PeproTech)) polarization. After 24 hours, cells were treated with 50 ng/mL of recombinant human IL-36γ (Bio-Techne) in complete medium. Cells were harvested after 6 or 24 hours. Cells were harvested at 6 hours for real-time qPCR to examine gene expression changes and at 24 hours for flow cytometry to assess surface marker expression.

### 2.2 Flow Cytometry

Macrophages were incubated with 10 μg/mL human Fc block (BD Biosciences) for 10 minutes at room temperature. Cells were then stained on ice for 30 minutes with fluorochrome-conjugated antibodies (Table S1) and for another 30-minute with LIVE/DEAD™ Fixable Near-IR Dead Cell Stain (Invitrogen). Cells were acquired using a CytoFLEX flow cytometer and the data was analysed using FlowJo (BD Biosciences, v10.9.0).

### 2.3 Real Time PCR and ELISA

Total mRNA was extracted from macrophages using the RNeasy Mini Kit (QIAGEN) and reverse-transcribed into cDNA with a High-Capacity cDNA Reverse Transcription Kit (Applied Biosystems). Real-time PCR was performed using the PowerUp™ SYBR™ Green Master Mix (Applied Biosystems) with primers for target and housekeeping (*GAPDH*) genes (sequences in Table S2). TNF-α and IL-8 were measured with the Human TNF-alpha Quantikine ELISA Kit (Bio-Techne) and Human IL-8 Quantikine ELISA Kit (Bio-Techne).

### 2.4 Measurement of Oxidized Low-Density Lipoproteins (oxLDL) Uptake

Human monocyte-derived macrophages were incubated at 37°C for 30 minutes in serum-free RPMI 1640-GlutaMAX™ containing 5 μg/mL DiI-conjugated human oxLDL (Thermo Fisher Scientific). oxLDL uptake was then quantified using a CytoFlex flow cytometer (Beckman Coulter), with detection of DiI fluorescence in the PE channel. A positive control was included by exposing cells to DiI-oxLDL for 24 hours. The data was analysed using FlowJo (BD Biosciences, v10.9.0).

### 2.5 Statistical Analysis

Comparisons between two independent groups were undertaken with Student’s unpaired t-test or the Mann-Whitney U test, as applicable. Comparisons between multiple independent groups were undertaken using one-way ANOVA with Tukey’s post-hoc test. The analysis of paired samples was implemented with Student’s paired t-test or Wilcoxon signed-rank test. When multiple paired comparisons were performed at the same time, a Bonferroni-Dunn correction was applied. P-values <0.05 were considered significant.

## 3. Results and Discussion

To recapitulate the functional extremes of macrophage plasticity, we differentiated human monocytes into pro-inflammatory (M1), tissue remodelling (M2a), and immune-regulatory (M2c) subsets (Fig. S1). We then assessed IL-36R expression on these cells. We found that the proportion of IL-36R–positive cells was higher in M2a (90.7%) compared to M2c (68.8%) and M1 (55.9%) macrophages (Fig. 1a). Consistently, IL-36R mean fluorescence intensity (MFI) was highest in M2a macrophages (Fig. 1a). This population may therefore be particularly sensitive to the effects of IL-36 cytokines.

**Fig.1.**
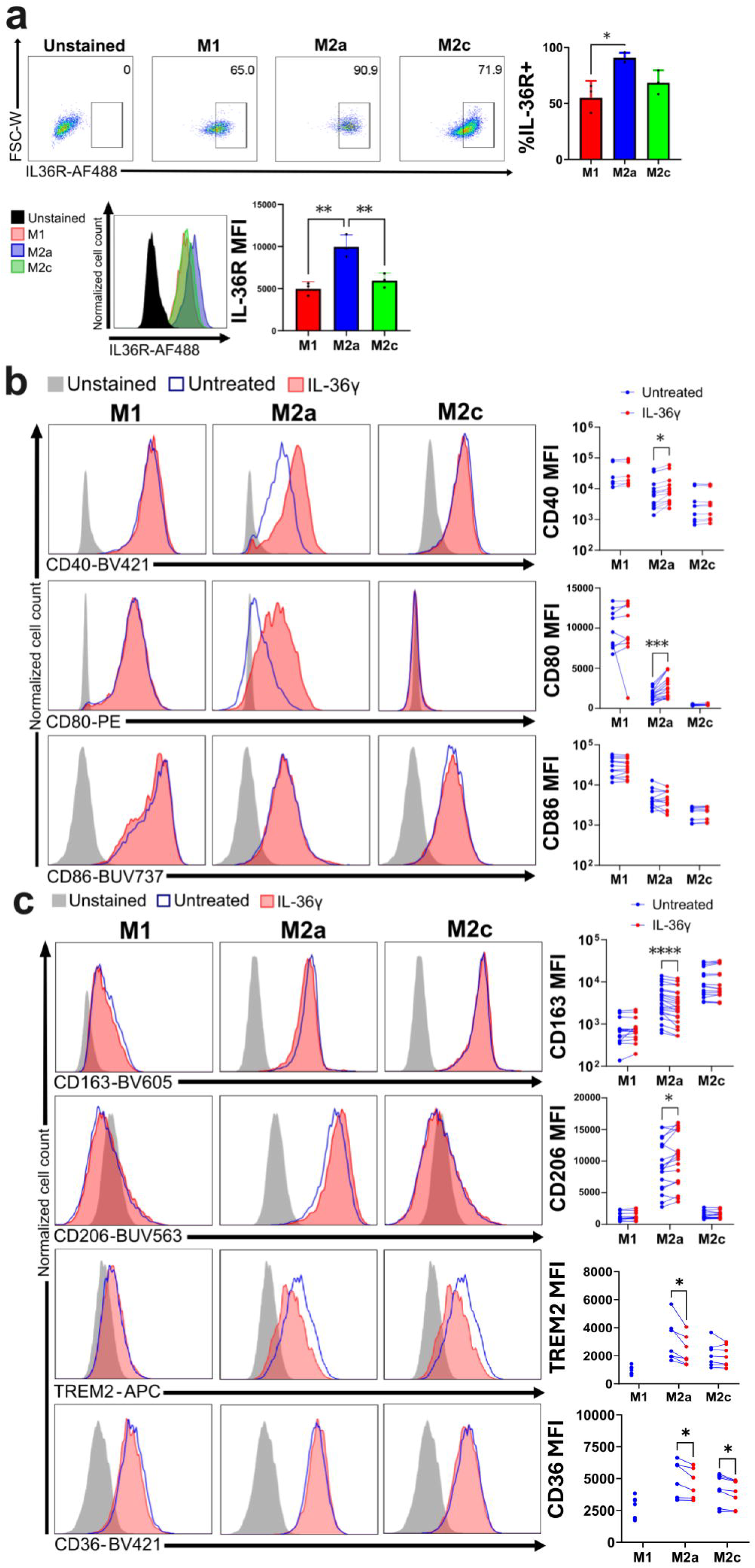
IL-36γ treatment shifts human M2a macrophages towards an M1 state. (a) (Left) Representative flow cytometry plots and bar plots showing the percentage of cells expressing IL-36R, gated on singlet live CD14+ cells. (Right) Representative flow cytometry histograms and bar plots showing the MFI of IL-36R in each subset. **P<0.01 (One-way ANOVA with Tukey’s post-hoc test). (b-c) Representative flow cytometry histograms (left) and paired dot plots (right) showing that IL-36γ treatment of monocyte-derived macrophages affects the expression of M1 (b) and M2 (c) markers *P<0.05, ***P<0.001, ****P<0.0001 (paired t-tests followed by Bonferroni-Dunn correction).

We next investigated how macrophage polarisation is affected by IL-36γ, the cytokine isoform that is most abundantly expressed in barrier tissues^2^. We specifically examined the effects IL-36γ on M2a and M2c macrophages, including the M1 subset as a reference for a proinflammatory state. We found that cytokine treatment had no significant effects on M2c cells. Conversely, we observed that in M2a macrophages, IL-36γ significantly up-regulated M1-associated molecules, including CD40 (1.34-fold, P<0.05) and CD80 (1.64-fold, P<0.001) (Fig. 1b). Meanwhile the analysis of M2 markers showed that IL-36γ induced modest increase in CD206 MFI (1.09-fold, P<0.05), but a pronounced reduction in CD163 expression (0.89-fold, P<0.0001) (Fig. 1c). To further investigate these findings, we examined the effects of IL-36γ on two additional proteins associated with M2 states, i.e. the TREM2 and CD36 lipid receptors^1^. We observed that both were significantly downregulated following IL-36γ treatment (TREM2: 0.76-fold, P<0.05; CD36: 0.92-fold, P<0.05) (Fig. 1c). Accordingly, we found that the uptake of oxidized low-density lipoprotein (a TREM2 and CD36 ligand) was significantly reduced in M2a macrophages that had been pre-treated with IL-36γ (0.90-fold, P<0.01) (Fig. 2a). Thus, IL-36γ upregulates M1 markers in M2a macrophages, while downregulating canonical and non-canonical M2 markers.

**Fig.2.**
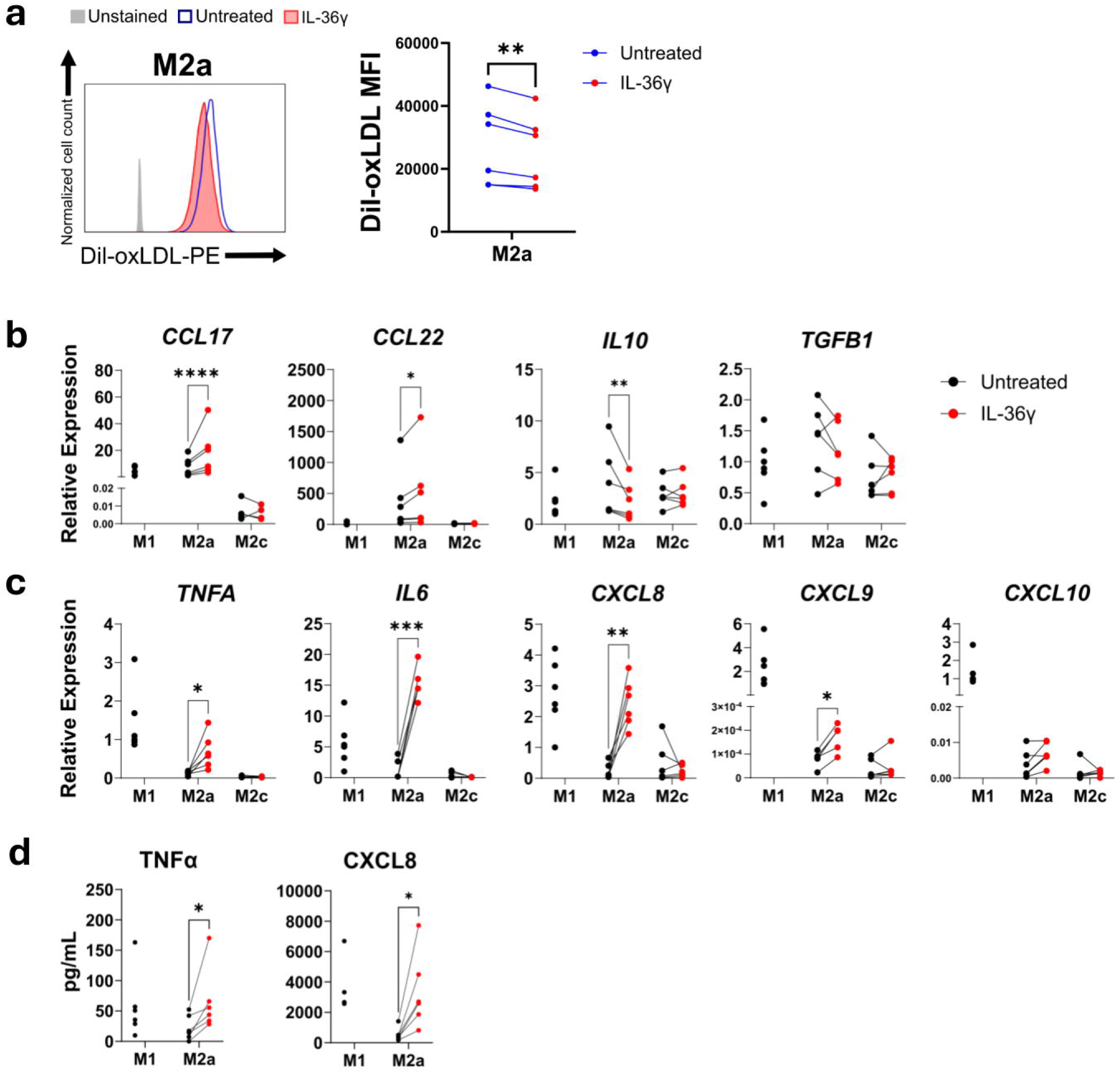
IL-36γ treatment affects lipid uptake and cytokine expression in human M2a macrophages. (a) Representative flow cytometry histogram (left) and paired dot plots (right) showing that IL-36γ treatment of M2a macrophages affects oxLDL uptake. (b-c) Paired dot plots showing the effect of IL-36γ on the mRNA expression of M2 (b) and M1 (c) cytokines. Gene expression was measured by real-time qPCR, using GAPDH as a housekeeping gene. (d) Paired dot plots showing the effect of IL-36γ on TNF-α and CXCL8 protein production. Data for untreated M1 macrophages is included in all plots as a reference for a pro-inflammatory state. *P<0.05, **P<0.01, ***P<0.001, ****P<0.0001 (paired t-test; a Bonferroni-Dunn correction was applied in b-d).

To further validate the phenotypic shift towards a pro-inflammatory state, we examined the effects of IL-36γ on M2 cytokine production. We found that IL-36γ treatment modestly up-regulated the mRNA expression of *CCL17* (2.4-fold; P<0.0001) and *CCL22* (1.4-fold; P<0.01), while downregulating *IL10* (1.8-fold, P<0.01) (Fig. 2b). Conversely, IL-36γ induced a substantial increase in the expression of M1 cytokine genes, including *TNFA* (5.0-fold; P<0.01), *IL6* (10.9-fold; P<0.0001), *CXCL8* (8.5-fold; P<0.01), *CXCL9* (2.1-fold; P<0.01) and *CXCL10* (1.8-fold; P<0.05) (Fig. 2c). Follow-up ELISAs of TNF and CXCL8 confirmed that IL-36γ increases the release of these two key M1 mediators (TNF: 3.0-fold, CXCL8 6.1-fold; P<0.05 for both) (Fig. 2d). Thus, the pro-inflammatory shift induced by IL-36γ treatment of M2a macrophages is also reflected in an increase production of M1 cytokines.

Earlier studies have demonstrated that IL-36 can activate NF-κB signalling and upregulate cytokine production in human macrophages^9,10^. Here, we build on these findings to systematically investigate the pro-inflammatory shifts induced by IL-36 treatment of human tissue-remodelling (M2a) and immune-regulatory (M2c) macrophages. We found that IL-36γ polarised M2a macrophages towards an M1-like phenotype. This was evident at the surface marker level, as IL-36γ upregulated the expression of CD40 and CD80, while downregulating that of CD163, TREM2, and CD36. The changes in cytokine production further supported this shift, with IL-36γ strongly inducing TNFα, IL-6, and CXCL8. In this context, the modest induction of CCL17 and CCL22 suggests that IL-36γ does not repolarise M2a macrophages but rather reshapes their profile towards a mixed state enriched in pro-inflammatory mediators.

We observed that the downregulation of TREM2 and CD36 was accompanied by a reduced lipoprotein (oxLDL) uptake. Of note, the lysosomal degradation of lipoproteins generates fatty acids, one of the main sources of energy for M2 macrophages^11^. Given that the inhibition of fatty acid oxidation impairs M2 functions^11^, it is tempting to speculate that the effects of IL-36γ on lipid uptake may contribute to the proinflammatory shifts induced by the cytokine.

Our study has some limitations. While we analysed the effects of IL-γ36 on critical surface markers and cytokines, we did not undertake transcriptomics studies that could have captured a broader spectrum of changes, including those affecting lipid metabolism genes. Likewise, we did not investigate how IL-36γ affects macrophage functions such as antigen presentation or T cell activation.

As our experiments were restricted to an in vitro system, future work should investigate the role of IL-36γ in shaping macrophage phenotypes in disease settings. In fact, excessive IL-36 activity has been implicated in inflammatory conditions (e.g. psoriasis, inflammatory bowel disease and pulmonary fibrosis) characterised by dysregulated macrophage polarization^2,12^. Conversely, IL-36 signalling has a protective role in some cancers (e.g. lung squamous cell carcinoma^13^) where the abundance of TREM2+ macrophages correlates with a poor prognosis^14^. Further studies should therefore investigate the potential of modulating IL-36γ signalling to reprogramme macrophage responses in different disease contexts.

## Author Contributions

FC, SNK and JYXS designed the study. JYXS performed the experiments with GO. JYXS and FC wrote the manuscript. FC and SNK supervised the study. All authors have reviewed and agreed to the final manuscript.

