## Supplementary Material for "IL-36γ signalling promotes a proinflammatory macrophage state that is associated with reduced lipid uptake"

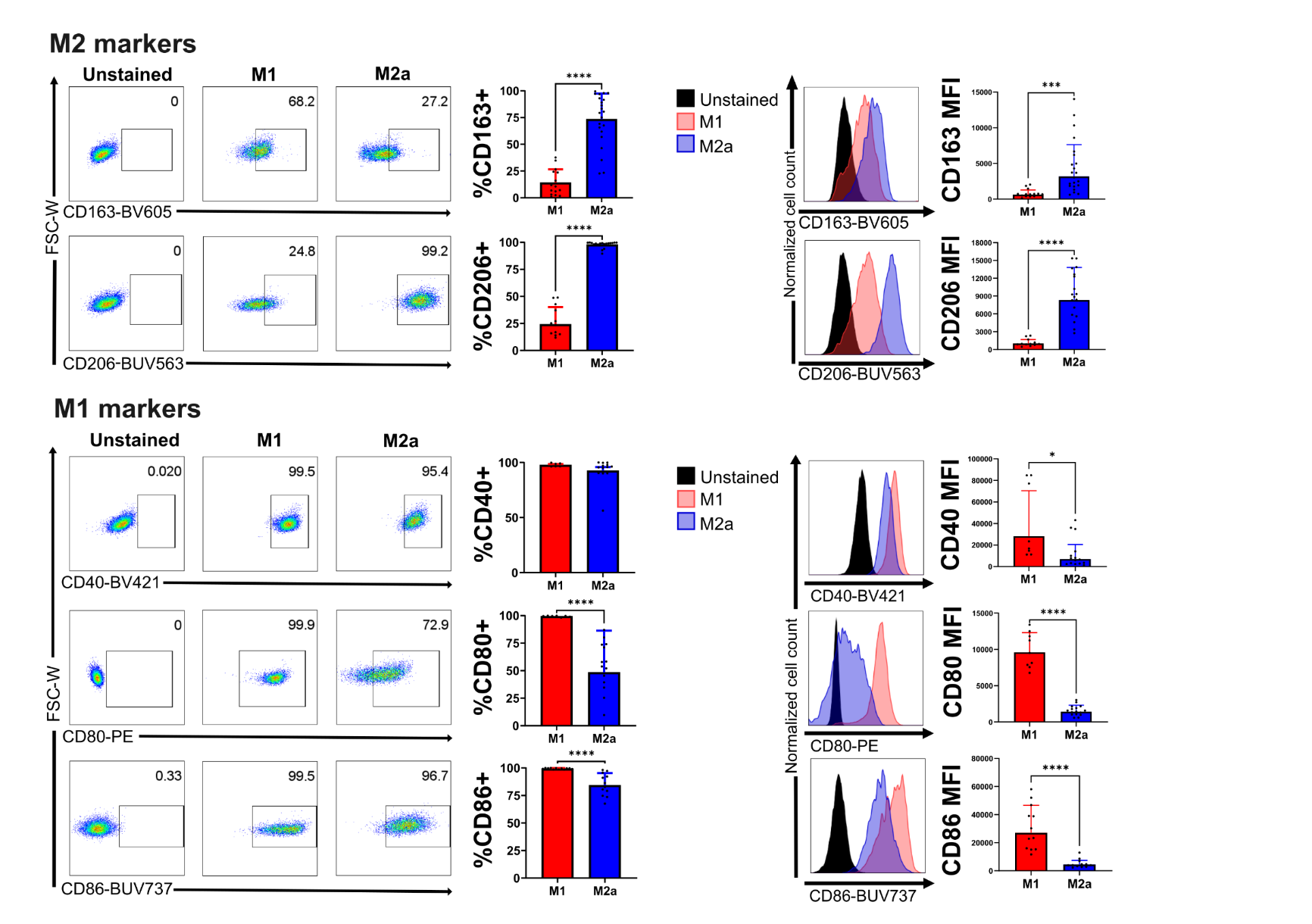

#### Figure S1: In vitro-polarised macrophage subsets express the expected surface markers. Left: Representative flow cytometry plots and bar plots showing the percentage of live CD14+ cells expressing each marker. Right: Representative flow cytometry histograms and bar plots showing the MFI of each marker in each subset. *P<0.05, ***P<0.001, ****P<0.0001 (Student’s t-tests).

### Supplementary Tables

**Table S1**: Antibody details

|  | **Supplier** | **Dilution** |
| --- | --- | --- |
| Monoclonal Mouse Anti-Human CD14-BUV395 [Clone: MφP9] | BD Biosciences | 1:250 |
| Monoclonal Mouse Anti-Human CD163-BV605 [Clone: GHI/61] | BioLegend | 1:70 |
| Monoclonal Mouse Anti-Human CD206-BUV563 [Clone: 19.2] | BD Biosciences | 1:130 |
| Monoclonal Mouse Anti-Human CD36-BV421 [Clone: 5-271] | BioLegend | 1:200 |
| Monoclonal Mouse Anti-Human CD80-PE [Clone: 2D10] | BioLegend | 1:34 |
| Monoclonal Mouse Anti-Human CD86-BUV737 [Clone: 2331 (FUN-1)] | BD Biosciences | 1:34 |
| Monoclonal Mouse Anti-Human CD40-BV421 [Clone: 5C3] | BioLegend | 1:34 |
| Monoclonal Rat Anti-Human/Mouse TREM2-APC [Clone: 237920] | R&D Systems | 1:200 |
| Polyclonal Rabbit Anti-Human IL1RL2 (IL-36R) Antibody^1^ | Fisher Scientific | 1:100 |
| Polyclonal Goat Anti-Rabbit IgG-AF488^1^ | Stratech Scientific | 1:200 |

^1^The anti-IL36R antibody is unconjugated, so fluorescence is detected following staining with a secondary goat anti-rabbit antibody

**Table S2**: Real-time PCR primers

| **Target** | **Forward (5'-3')** | **Reverse (5'-3')** | **Product (bp)** |
| --- | --- | --- | --- |
| ***CCL17*** | ACTTCAAGGGAGCCATTCCC | CCCTGCACAGTTACAAAAACGA | 97 |
| ***CCL22*** | CTGCGCGTGGTGAAACACTT | TCCCTGAAGGTTAGCAACACCA | 77 |
| ***CXCL8*** | GAGAAGTTTTTGAAGAGGGCTGA | CTTCACTGATTCTTGGATACCACA | 71 |
| ***CXCL9*** | AAGTGGTGTTCTTTTCCTCTTGG | CACTACTGGGGTTCCTTGCAC | 70 |
| ***CXCL10*** | AAGCAGTTAGCAAGGAAAGGTCTA | GCAGCCTCTGTGTGGTCCAT | 96 |
| ***GAPDH*** | CGGAGTCAACGGATTTGGTC | AATGAAGGGGTCATTGATGGCA | 97 |
| ***IL10*** | TTGCTGGAGGACTTTAAGGGTT | GTTCTCAGCTTGGGGCATCA | 93 |
| ***IL6*** | CCTTCTCCACAAGCGCCTTC | AAGGCAGCAGGCAACACCA | 70 |
| ***TGFB1*** | ACGTGGAGCTGTACCAGAAAT | AGATAACCACTCTGGCGAGTC | 92 |
| ***TNFA*** | CTCTCTCTAATCAGCCCTCTGG | GCTTGAGGGTTTGCTACAACA | 95 |
